# Assessing Perceptual Chromatic Equiluminance Using a Reflexive Pupillary Response

**DOI:** 10.1101/2023.10.19.563116

**Authors:** Liu Ye, Bridget W. Mahony, Xiaochun Wang, Pierre Daye, Wei Wang, Patrick Cavanagh, Pierre Pouget, Ian Max Andolina

## Abstract

Equiluminant stimuli help assess the integrity of colour perception and the relationship of colour to other visual features. As a result of individual variation, it is necessary to calibrate experimental visual stimuli to suit each individual’s unique equiluminant ratio. Most traditional methods rely on training observers to report their subjective equiluminance point. Such paradigms cannot easily be implemented on pre-verbal or non-verbal observers. Here, we present a novel Pupil Frequency-Tagging Method (PFTM) for detecting a participant’s unique equiluminance point without verbal instruction and with minimal training. PFTM analyses reflexive pupil oscillations induced by slow (< 2 Hz) temporal alternations between coloured stimuli. Two equiluminant stimuli will induce a similar pupil dilation response regardless of colour; therefore, an observer’s equiluminant point can be identified as the luminance ratio between two colours for which the oscillatory amplitude of the pupil at the tagged frequency is minimal. We compared pupillometry-based equiluminance ratios to those obtained with two established techniques in humans: minimum flicker and minimum motion. In addition, we estimated the equiluminance point in non-human primates, demonstrating that this new technique can be successfully employed in non-verbal subjects.

**Key Points:**

- Measuring equiluminance is challenging in non-verbal subjects.
- The pupil diameter varies as perceived brightness changes.
- Utilising oscillatory pupillometry, PFTM provides a way to measure equiluminance points without training or the need for a subjective report.

## 1. Introduction

Perception of the chromatic properties of light is essential for adaptive visual behaviours in many animals^1^, including human and non-human primates^2^. While most eutherian mammals are dichromats, about 30-40 million years ago, a mutation of the X chromosome-linked gene for the long wavelength-sensitive cone pigment resulted in trichromatic vision in catarrhine primates (the family that includes old-world monkeys, apes, and humans). Substantial individual variation in cone distributions and the subsequent processing across the visual cortex inevitably leads to substantial subjective differences in the perceptual properties of colour^3,4^.

One such property that differs among individuals is the response of the luminance pathways to different chromatic inputs. This response is proportional to a stimulus’ objective luminance as defined by luminous intensity per unit area (candelas per square meter, cd/m^2^), but the proportion varies across individuals. In experiments of colour perception, it is essential to set the colours of the stimuli to subjectively equal luminance levels to better isolate the visual system’s elements sensitive to colour. For example, equiluminant colour stimuli optimally isolate midget cells’ relative contribution to the parvocellular layers of the lateral geniculate nucleus (sensitive to chromatic contrast) in constructing visual perception5,6. However, there is substantial individual variation in the luminance response to stimuli of the same objective luminance value^7^. As a result, an individual’s unique equiluminance ratio must be determined in order to minimise response in the observer’s luminance pathway. While there are several established methods of evaluating equiluminance ratios^8^, most require the cooperation and understanding of the individual being tested. They are less effective for assessing the equiluminance ratio of nonverbal or preverbal populations, such as non-human primates or human infants^9^. Optokinetic nystagmus (OKN) has been used to measure the equiluminance point in human infants, amphibians, and fish^1–12^. However, the OKN reflex critically depends on the processing of motion, which has a dominant input from magnocellular pathways that provide weak contributions to colour pathways^13,14^. Additionally, because similar oculomotor structures control the OKN and ocular fixation, the OKN can be differentially suppressed via the frontal eye fields during attentive fixation in human and non-human primates^15,16^. This voluntary control of the OKN response makes its use problematic for populations such as non-human primates with extensive training in fixation.

There is a growing appreciation of how, in addition to classical reflexive mechanisms, subjective cognitive processing also modifies pupil diameter^17,18^. Perhaps the clearest example of this comes from studies showing that learned categories (i.e. “the Sun is very bright”) ^19–21^ or illusions of brightness in humans^22^, and even rodents^23^, can change pupil size. Additionally, there is considerable evidence that the pupillary response is sensitive to inputs from the extrastriate colour pathways in human and non-human primates^24–27^. While the pupillary response to colour can be impacted by attention^28^, this is not the global antagonistic impact it has on the OKN reflex. Therefore, we developed a new method of measuring equiluminance ratios that provides an objective measure in verbal and nonverbal populations. This uses a low-cost eye tracking device along with rapid and straightforward signal processing methods. Pupil diameter is mainly driven by perceived luminance independently of colour^22^ so that a flickering stimulus that alternates between two colours will cause the pupil diameter to oscillate at the same frequency as the luminance variation in this stimulus (hereto forth, the “tagging frequency”). With this Pupil Frequency-Tagging Method (PFTM) the amplitude of the oscillations at the tagging frequency will be proportional to the perceived luminance difference between the two colours — as long as the frequency is slow enough to drive a reliable pupillary response (< 2 Hz)^29^ and fast enough to minimise input from the melanopsin pathway that also drives the pupil response for prolonged changes (< 1 Hz)^30,31^ (noting that the melanopsin response has little to no impact on colour perception^32^). The pupil oscillations will be at a minimum when the two alternating colours have matched perceived luminance. Therefore, an observer’s unique equiluminance point can be identified as the luminance ratio between two colours at which the Fast Fourier Transform (FFT) amplitude at the tagging frequency is minimal.

## 2. Results

### 2.1. Exemplar PFTM Measurements in Human

We first sought to measure whether the PFTM stimulus successfully drove oscillations in pupil diameter that were synchronous with the frequency of the alternations of two different luminances of the visual stimuli. The stimulus itself comprised a 20° smoothed central disc that changed between two stimuli (either two luminance values of the same colour, or two luminance values of two different colours) every 0.283 seconds, presented against a physically equiluminant grey background (**Figure 1**a). As the participant maintained fixation, pupil diameter was recorded with an eye-tracker with sample times aligned to stimulus onset. **Figure 1**b shows an example of the raw pupil diameter plotted against time: the main features of the pupillary stimulus-locked oscillation that vary with luminance are a *mean baseline change*, a *variable oscillatory amplitude* and a *phase reversal* for luminances above and below a minimal amplitude.

**Figure 1:**
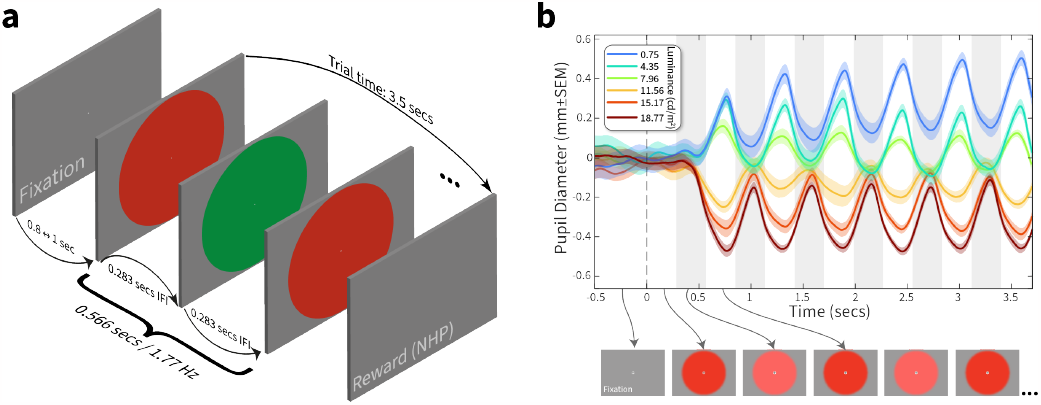
PFTM stimulus design. (**a**) Participants initiated a trial by fixating on a central spot for 0.5 – 1 seconds before a 20° disc oscillated between fixed and variable luminance colours every 0.283 seconds (34 frames at 120 Hz display refresh) resulting in a tagging frequency of 1.77 Hz. The trial continued for 3.5 seconds as participants maintained their fixation. (**b**) the exemplar plot of relative pupil diameter averaged over ten randomised trials (±SEM); each curve colour represents a different variable luminance value shown in the legend. Alternating grey shading represents the 0.283 second duration of the fixed and variable colours (illustrated by arrowed example frames below the axis).

We first used the same colour (eg. a fixed red and variable red) to verify that the pupillary oscillation showed a minimum at the fixed luminance. In all cases we found that the oscillation in the raw pupil diameter disappeared only when the fixed and variable colour were identical. We next wanted to test how the oscillation varied when testing a different fixed and variable colour. A full example of the analysis necessary to quantify a perceptual minimum between two different colours are shown in **Figure 2** through **Figure 4**. They demonstrate pupil diameter oscillations, amplitude extracted via the FFT transform, and normalised amplitude at the tagging frequency plotted across all variable luminances, respectively. For this human exemplar (participant 2), we held red fixed at 21.72 cd/ m^2^, and varied green in ten equally spaced steps between 0 cd/m^2^ and 41.2 cd/m^2^ (**Figure 2a**). We chose 5 repeated trials of each luminance to minimise task duration, leading to a total task time of 5 minutes and 25 seconds.

**Figure 2:**
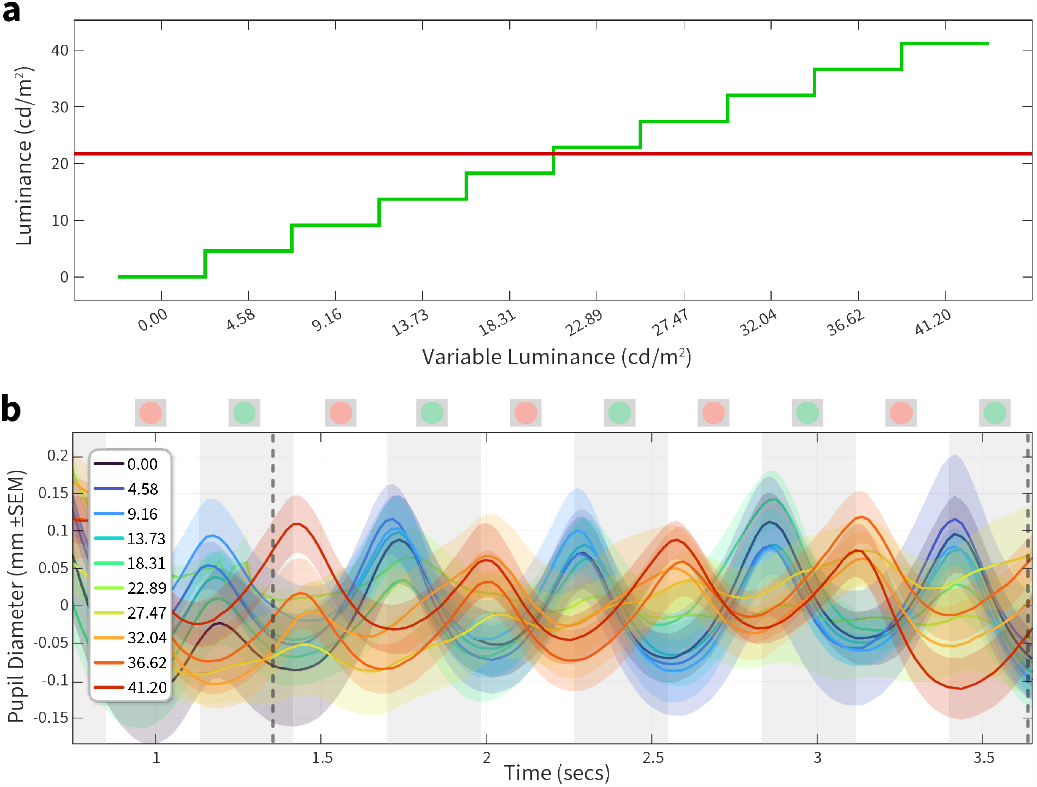
stimulus values and resultant pupil diameter measurements. (**a**) Schematic of the luminance steps between a fixed red (21.72 cd/m^2^) against 10 steps of green ranging from 0 to 41.2 cd/ m^2^. (**b**) Mean-corrected pupil diameter is plotted against time; the white and grey vertical bands show the exchanges of the two luminances with a frequency of 1.77 Hz (schematics of the stimulus shown above the axis). The colour of each curve represents the luminance steps shown in (a). Each luminance was repeated five times and data is plotted ±SEM Dotted vertical lines indicate the start and end points where the FFT amplitude was computed in **Figure 3**.

**Figure 3:**
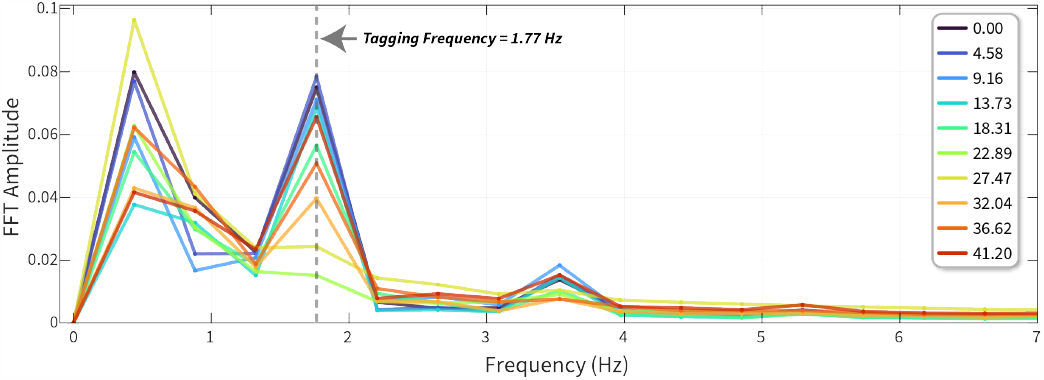
FFT analysis of pupil oscillation. FFT amplitude between 0 _↔_ 7 Hz is plotted for each variable luminance condition. The amplitude at 0 Hz, equivalent to the mean, is not seen here as the pupil oscillations were first baseline-corrected using the mean pupil diameter. The dotted vertical line indicates the tagging frequency (1.77 Hz) at which the stimulus oscillated.

**Figure 4:**
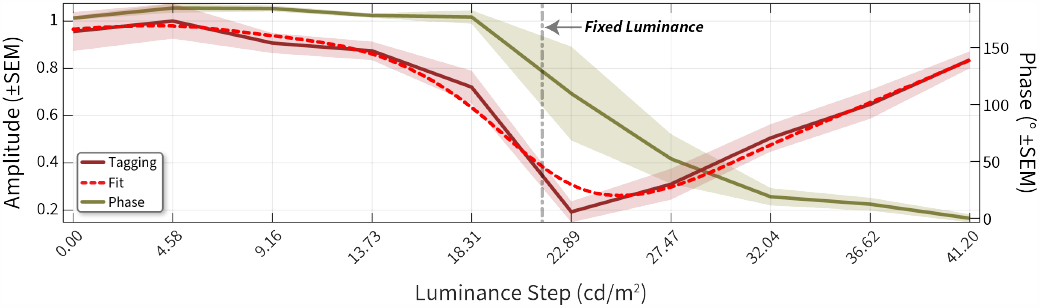
normalised FFT amplitude. amplitude (leftward Y-axis) at the tagging frequency is plotted across luminance conditions (X-axis). Vertical grey dotted line denotes the fixed luminance value. We used a smoothing spline function (dotted red line) fit to the amplitude curve and estimated the ratio of red:green from this fit. The green curve plots the phase (rightward Y-axis) at the tagging frequency plotted across luminance conditions. Shading ±1SEM.

The stimulus oscillation is illustrated by alternating grey bands and stimulus icons in **Figure 2b**. Pupil changes are slow, and so we excluded the first second after stimulus onset then used the mean amplitude of the measurement range to correct each conditions response before averaging. The critical features of the pupil oscillation were the minimum in amplitude as a function of the variable luminance and a clear reversal in the oscillatory phase, with an almost complete 180° inversion of phase between the darkest and lightest values of the variable test colours.

The FFT can extract amplitude and phase measurements from a time–series signal. We first plotted the FFT amplitudes across all frequencies ranging from 0 Hz to 7 Hz. 0 Hz represents the mean baseline change, and in this case, we baseline-corrected the pupil diameter signals resulting in no values at 0 Hz. The tagging frequency was 1.77 Hz, and we can see an amplitude peak at that frequency whose value changes across luminance. The amplitudes (normalised to the largest amplitude) for each luminance at the tagging frequency are shown in **Figure 4**. We predicted that the amplitude of pupillary oscillations should be minimised when the two stimuli had a similar luminance.

We noted a clear change in normalised amplitude with luminance, and the minimum was observed at 22.89 cd/m^2^ of green (red luminance denoted by dotted line). This denoted a clear shift from the physical equiluminant point. We also noted that the phase transition started after 18.31 and ended at 32.04. The best-fitting spline had its minimum more closely aligned to the mid-point of the phase transition, and we therefore used the fitted curve to estimate the minimum ratio, in this participant giving a value of 0.79.

### 2.2. Comparisons of PFTM with Flicker and Motion Tasks in Humans

Previous methods that estimate a participant’s equiluminance point include tasks in which they must subjectively decide (using either an alternate forced choice or a matching paradigm) on a stimulus minimum.

For example, in a heterochromatic minimum flicker task (**Figure 5a**), a participant can change the luminance of one test colour as a square wave reverses colour contrast over time to minimise the perception of flicker. In a minimum motion task, the participant can choose the direction of motion (for example, using a leftwards or rightwards 2-alternate forced choice [2–AFC]) and the point at which leftward and rightward choices occur equally is taken to be the perceptual minimum (**Figure 5b**). We used a matching paradigm for our minimum flicker and a randomised 2–AFC task for minimum motion.

**Figure 5:**
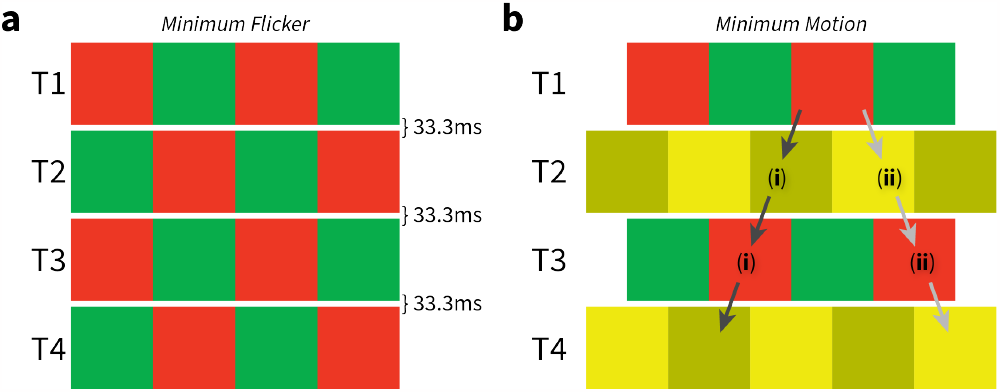
subjective equiluminance testing. (**a**) Minimum flicker: a red/green square wave grating alternates 30 times per second with a phase-reversed version that replaces it at the same location. With the luminance of the red bars fixed, the participant adjusts the relative luminance of green bars to reduce the amount of flicker seen in the display. (**b**) Minimum motion: coloured gratings are shown in a repeating sequence, at times T1 to T4. Each grating shifts sideways by one-quarter cycle (half of a bar width) from its predecessor. The direction of the parent motion, shown by the arrows, depends on the relative luminance of the red and green bars. (**i**) When the red bars are darker than the green bars, the red bars in the grating at time T1 (or T3) appear to jump leftward to the dark yellow bars in the grating at time T2 (or T4). (**ii**) Conversely, when the red bars are lighter than the green bars, the red bars appear to jump rightward to the light-yellow bars. The luminance contrast between the light and dark yellow bars was 40%.

The minimum-flicker, minimum-motion and PFTM settings for red versus green for the eight participants are shown in the three columns of **Table 1**, respectively. We visualised these values using a raincloud plot (**Figure 6a**) and as a correlation matrix (**Figure 6b**). Overall, no significant differences existed between any group means (Flicker:Motion p = 0.69 / BF_10_ = 0.36; Flicker:PFTM p = 0.46 / BF_10_ = 0.95; Motion:PFTM p = 0.36 / BF_10_ = 0.7; Holm post-hoc corrected repeated-measures ANOVA / Bayesian paired t-tests).

**Table 1:** red to green ratios at equiluminance comparing individual participant values for minimum flicker, minimum motion and PFTM (mean ± SEM).

| Participant | Flicker | Motion | PFTM |
| --- | --- | --- | --- |
| 1 | 0.83 | 0.84 | 0.89 |
| 2 | 0.81 | 0.94 | 0.79 |
| 3 | 0.93 | 0.87 | 0.9 |
| 4 | 0.88 | 0.94 | 0.82 |
| 5 | 0.86 | 0.84 | 0.88 |
| 6 | 0.84 | 0.79 | 0.74 |
| 7 | 1.01 | 1.09 | 0.96 |
| 8 | 0.86 | 0.79 | 0.79 |
| Mean±SEM | 0.88 ± 0.02 | 0.89 ± 0.04 | 0.85 ± 0.03 |

**Figure 6:**
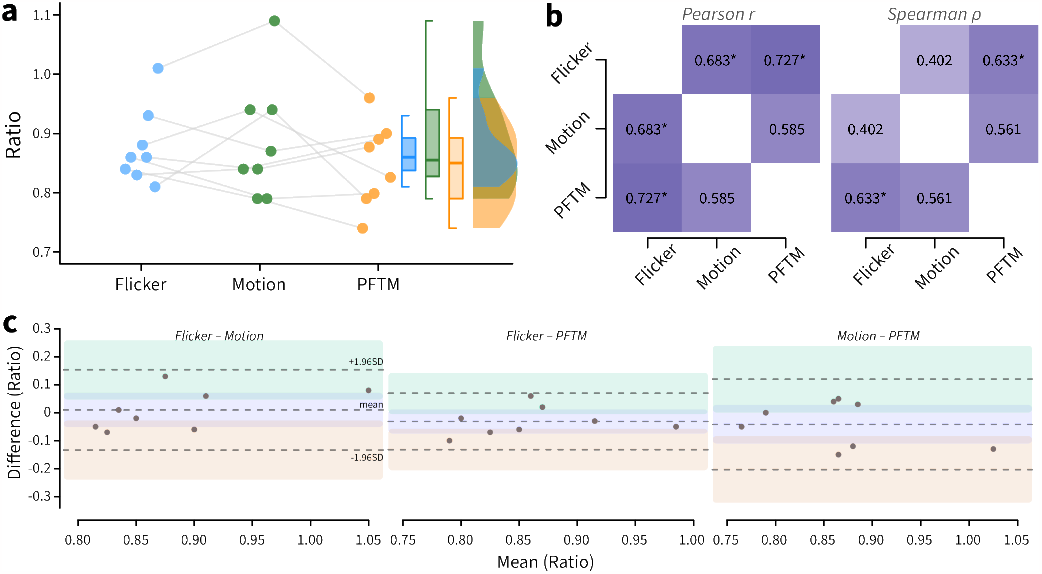
comparison of minimum flicker, minimum motion and PFTM paradigms for 8 human observers. (**a**) The raw values of the red-green ratio are plotted for PFTM (right), minimum motion (middle) and minimum flicker paradigms (left). The paired raincloud plots also show box and violin density plots across the 8 participants. (**b**) Pearson r and Spearman ρ correlation matrices across the three different tasks. * = significant positive correlation at p < 0.05. (**c**) Bland-Altman pair-wise reliability plots, dotted lines denote the mean ±1.96 SD; shaded areas are the 95% confidence intervals of the mean and upper/lower SD.

Testing for bivariate normality suggested that only Flicker:Motion violated the assumption of normality requiring the use of a nonlinear correlation test (Shapiro-Wilk = 0.789, p = 0.022). The correlation between Flicker:Motion was weakly positively correlated without reaching significance (one-tailed Spearman ρ = 0.402, effect size = 0.427, p = 0.161). We found that Flicker:PFTM exhibited the strongest significant positive correlation (one-tailed Pearson r = 0.727, effect size = 0.922, p = 0.021), and Motion:PFTM showed a moderate positive correlation that did not reach significance (one-tailed Pearson r = 0.585, effect size = 0.67, p = 0.064). We also performed pairwise reliability plots (**Figure 6c**) that suggested that there are no substantial differences in agreement across the three methods (Flicker:Motion −0.13 ≺ 0.01 ≻ 0.15 | Flicker:PFTM −0.13≺-0.03 ≻ 0.07 | Motion:PFTM −0.2 ≺ −0.04 ≻ 0.12 | −1.96 SD ≺ mean ≻ 1.96 SD). Overall, the values were within the previously reported range for colour normal observers, and the weak correlation between minimum flicker and motion has been previously described^33^,8.

### 2.3. Measuring PFTM in Non-human Primates

It is often optimal for non-verbal subjects like children or non-human primates for test sessions to contain independent trials, where positive feedback and encouragement (in children) and fruit or food rewards (non-human primates) can be assigned for correct behaviour. Previous studies have shown that pupil responses to colour in human and non-human primates exhibit similar properties^2^5. To test this, we used our PFTM protocol with four non-human primates that had previously been trained to keep their eyes steady. They were required to fixate a central spot for 3.5 seconds while the stimulus oscillated between the fixed and a variable luminance at the tagging frequency. We first used the same colour to test that the PFTM could identify a clear minimum. Given our subject’s daily trial completion rates, we increased the sampled luminance density and used a greater number of repeated trials at each luminance.

We first sought to confirm that the PFTM method could drive robust pupil signals with features similar to those observed in humans using a same–colour test. **Figure 7a** plots control data for the condition where both the fixed and variable luminance were in the green channel. The fixed luminance was set at 11.04 cd/m^2^, and 21 variable values were linearly spaced between 1.6 cd/m^2^ and 33.05 cd/ m^2^. Ten repeated trials were taken for each variable luminance value (210 trials total), and all trial placements were fully randomised and interleaved. We baseline-corrected using the pre-stimulus mean amplitude (around 0 seconds) to visualise any potential baseline changes during visual stimulation. After an initial response latency, the pupil diameter increases (signifying dilation caused by a lower perceived brightness) or decreases (signifying constriction caused by a higher perceived brightness) with a baseline offset positively correlated with the difference between the fixed and variable values. After about 0.7 seconds, the signal oscillated with a period matched to the stimulus reversals. In addition, the phase of the signal peaks for variable values lower than 11.04 cd/m^2^ reverses for variable values higher than 11.04 cd/m^2^. This confirmed that the pupil signals in non-human primate had the same features as those observed in humans.

**Figure 7:**
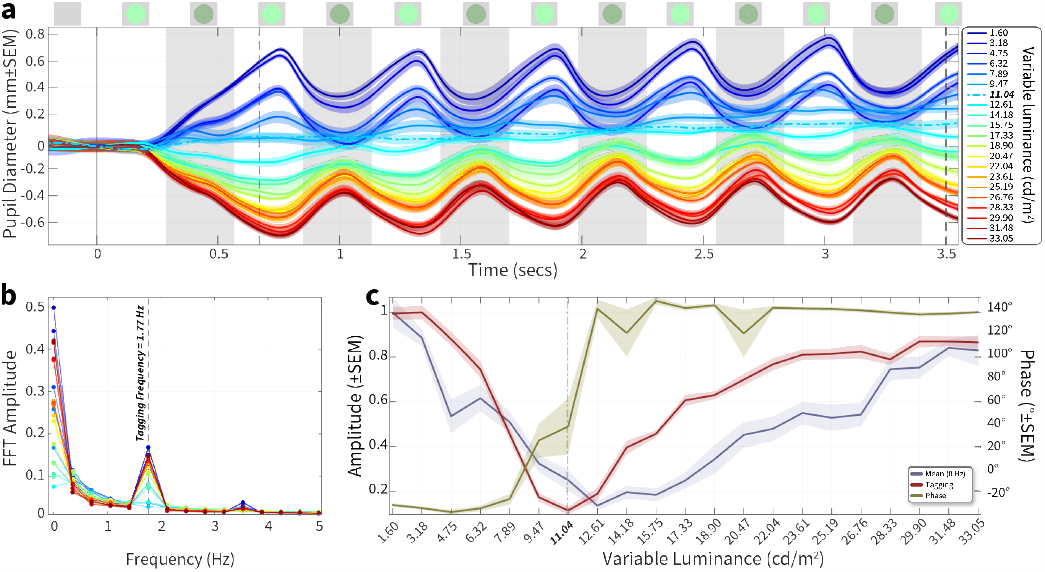
PFTM same-colour session in a non-human primate. (**a**) pupil diameter plotted against time with a fixed luminance of 11.04 cd/m^2^; plot colour of each curve represents the varying luminance condition (10 repeated trials) plotted ±SEM Each curve is baseline corrected using the mean amplitude ±0.2 around 0 seconds. (**b**) FFT amplitude between 0_↔_5 Hz is plotted for each variable condition; 0 Hz is equivalent to the mean, and the dotted vertical line indicates the tagging frequency at which the stimulus oscillated. (**c**) Normalised FFT amplitude (leftward Y-axis) at the mean (blue) and tagging frequency (red) plotted across luminance conditions (X-axis). The green curve plots the phase (rightward Y-axis) at the tagging frequency plotted across luminance.

To quantify this, we computed the FFT and focused on the frequencies at the tagging frequency (**Figure 7b**). As we did not remove the baseline offset, there is substantial signal at 0 Hz, which demonstrates the mean offset from the baseline, and the largest values are those with the greatest differential to the fixed value. There is also a clear peak at the tagging frequency, whose magnitude is again positively correlated with the difference between fixed and variable values. To better measure these relationships, we replotted the FFT amplitude data for the mean (0 Hz) and tagging frequency against variable stimulus luminance (**Figure 7c**). There is a clear minimum at the equiluminant 11.04 cd/m^2^ point for the amplitude extracted at the tagging frequency (dark red curve). The curve plotted for the amplitude extracted for the mean (0 Hz dark blue) also shows a minimum; however, it is at 12.61 cd/m^2^, missing the equiluminant point, and exhibits a somewhat broader tuning relationship across luminance (it was common that the mean amplitude was not a reliable indicator of an equiluminant point). Finally, the phase calculated at the tagging frequency shows the reversal identified in **Figure 7a**, going from around −30° at luminances lower than, to around 140° for luminances higher than the equiluminant point.

We next held red as the fixed colour and varied green around this point (as in the human experiments) to test our four participants. We found clear minima that differed from the physical luminance (CIE) matches (**Table 2**). The ratio values we observed were qualitatively within the range of that seen for the human participants.

**Table 2:** ratios of red to green for four rhesus macaques for minimum amplitude at the tagging frequency. The minimum was were defined as the lowest amplitude at the tagging frequency and confirmed by a phase inversion across the minimum.

|  | Aman | Fanfan | Rhaegel | Wallace |
| --- | --- | --- | --- | --- |
| Ratio | 0.89 | 0.93 | 1.11 | 0.9 |

Our non-human participants could maintain fixation for 3 to 4 seconds, but not all participants may be able to maintain fixation for this length of time. We were therefore interested in seeing whether a shorter data collection can yield a measurable minimum using a post-hoc analysis. **Figure 8** shows the pupil data for participant Fanfan comparing the raw data (upper) and FFT amplitude and phase (lower) for a single cycle (**Figure 8a**) or 5 cycles (**Figure 8b**).

**Figure 8:**
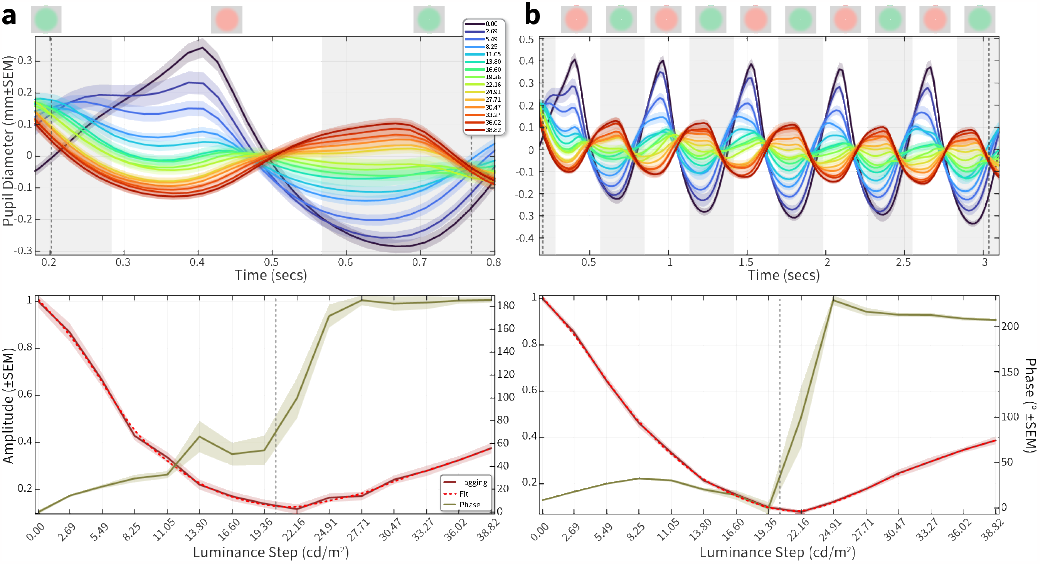
impact of data length on PFTM measurement in a non-human participant. (**a**) upper plot: the average (20 repeated trials) mean-corrected pupil diameter for one cycle (upper plot) across 15 different green luminance values for a fixed red of 20.34 cd/m^2^ (dotted grey lines represent the time-period from which the FFT was calculated). lower plot: the resultant amplitude (red, left Y-axis) and phase (green, right Y-axis) across 15 green luminance values, with a fitted (dashed red line) minimum yielding a ratio of 0.94 (dotted grey line represents physical luminance of red) (**b**) Average pupil diameter for five cycles (including the cycle from (**a**) and the resultant amplitude and phase across 15 green luminance values, with a fitted (dashed red line) minima yielding a ratio of 0.93; note the smaller error bars and more precise phase transition, but little change in the overall RG ratio.

Although the error bars are larger for the single cycle and phase transition more variable, the overall minimum defined by amplitude at the tagging frequency is preserved. We can therefore trade-off fixation length against noise depending on the participant’s ability to maintain fixation. We also tested downsampling the raw pupil data to 120 Hz, considering eye-trackers with a lower sampling rates, which had no impact on the FFT results (data not shown).

## 3. Discussion

We successfully identified the equiluminance point of participants by analysing pupil oscillations at the tagging frequency (1.77 Hz) and compared these values to measurements obtained using traditional photometric tasks. Our results show that the amplitude of pupil dilation oscillations in the tagging frequency band closely corresponds to the difference in luminance values when a single colour was used in the flickering stimuli and that these luminance-induced pupil responses are reduced at equiluminance when two colours are used. This measurement was significantly correlated to a previously well–established minimum flicker measurement. Consequently, PFTM enables measurement of equiluminance at colour exchange rates far below the higher temporal rates used in the flicker and motion tasks. Pupillary Frequency-Tagging has a significant advantage over established subjective techniques by objectively measuring an obligatory, automated response (pupil dilation) rather than relying on subjective responses about the appearance of the stimulus.

The pattern of ratio values for the two subjective methods compared to the PFTM shown in **Figure 6** show fair, but not exact correlation. Two accounts of these differences are worth discussing. First, it is important to note that the stimuli used in the PFTM, the minimum flicker, and minimum motion tests differ in several respects. Firstly, the temporal frequencies used are different across the three tasks. Changes in temporal frequency can profoundly affect neuronal responses in the retina and at cortical stages, which could thus account for some of the observed differences. Second, and perhaps more importantly, the circuits underlying the autonomic pupillary response and subjective responses differ. The “fast” circuit involved in the pupil constriction response — from the retina to the iris sphincter bypasses the thalamocortical pathway. This is not the case with subjective pupil responses that reflect the processing of downstream cortical neurons along the visual processing pathways from V1 to V4 and beyond^24,25,34,35,27^. At these higher levels of processing, neurons have an exquisite sensitivity to colour contrast, and colour/luminance perception results from complex interactions that initiate within the magnocellular, parvocellular and koniocellular cells from V1 onwards^36^. These distinctions in the relative contributions of the neural pathways may account for the some of the differences found here. However, it is known that feedback from frontal and occipital cortices via the locus coeruleus (LC), superior colliculus (SC), and hypothalamic orexin neurons do ultimately target the same nuceli^37,38,18,39^. It is also worth noting that the Spearman correlation between minimum motion and minimum flicker (both subjective methods) were the lowest among the three methods (**Figure 6**), therefore any such anatomical pathway differences cannot fully explain differences in the perceptual equiluminant point between methods.

Equiluminant stimuli are often used to reduce the relative contributions of magnocellular cells to visual processing to study the contributions of the parvocellular cells “in isolation” (though noting the contested effects on parvo vs. magno pathways^40^). A critical question arises concerning the best strategy to do so. At first, reducing the overall contribution of magnocellular cells at the retinal level may seem optimal, as cortical processing builds upon retinal inputs. One should be cautious however, in part because this view relies on underlying assumptions concerned with the projections of RGCs to the pupillary circuits that differ from their projections to the cortex. In this respect, recent detailed publications^41,42^ reveal the complex interactions within the retina before projecting to the pretectal olivary nucleus (PON) and Edinger-Westphal nucleus (EWN, that project to the ciliary ganglion driving the iris sphincter). The dominant view is that intrinsically photosensitive retinal ganglion cells (ipRGCs) containing melanopsin^43^ are the dominant drive to pupillary circuits for stimuli with temporal frequencies at 1 Hz or lower^30^. Our stimuli were modulated at 1.77 Hz. Interactions of ipRGCs with rods, cones, and RGCs have been consistently reported and can account for the pupillary responses to coloured stimuli not directly stimulating ipRGCs (as the absorption spectrum of melanopsin lies in the blue range — short wavelength) at lower temporal rates. Although it is out of the scope of this report to describe these circuits in detail, one should keep in mind that dedicated circuits exist to drive pupillary responses that differ from those driving cortical neurons through the thalamocortical pathways.

We used a method-of-constants presentation method for this study, but preliminary experiments also tested continual stimulation^44^ with no trial-by-trial structure. Subjects continued fixation and could blink etc. while the variable luminance ramped from a minima to maxima. By using post-hoc signal processing, we could also successfully extract the equiluminant ratio (data not shown). An alternative paradigm could use an adaptive staircase^45^ and given the computational simplicity to run the FFT in real time an adaptive staircase driven by the FFT amplitude of the previous trial could significantly reduce the time it takes to find the equiluminant point. Likewise, a heuristic that uses the phase–transition to objectively weight the minimum–amplitude should converge more quickly on the minimum than a fitted curve alone. Measurements using PFTM should extend to blue and yellow, and the use of alternative colour spaces. Finally, we expect that exploring the spatial extent of the PFTM stimulus, given the known fall-off of colour-sensitive cells in the retina, could further optimise the PFTM technique in future experiments.

A primary predicted application of the Pupil Frequency-Tagging Method described here is as a tool to assess colour blindness based on equiluminance settings^7^. It is equally as applicable to cooperative human adults as to infants and non-human animal subjects, improving previous colour blindness detection techniques and providing an opportunity for novel research trajectories. Additionally, PFTM is helpful as a method to validate that stimuli used in studies of colour perception are truly equiluminant. This is a critical variable to control for, as equiluminant stimuli ensure that the effects of colour perception are better isolated from those of luminance. For example, dominant theories suggest that the saccadic inhibition (SI) effect, observed when a visual mask or “distractor” placed over a saccadic target consequently inhibits saccadic movements, is driven by superior collicular pathways^46^,^47^. While the SC and its afferent pathways were not thought to represent colour stimuli directly^48^–^50^, recent studies in non-human primates using physically isoluminant colour change detection does exhibit SC activity in non-human primates (which they speculated to come from V4 or IT^51^,^52^). In both this task and the SI task, it would be beneficial that the colours used were perceptually matched to ensure that the visual circuit cannot identify the target and produce saccades or modify behaviour based on residual luminance signals alone. Studies that test neurodevelopmental changes of perception in human or non-human infants are another ideal target^53^, and PFTM would be a valid method to assess the infants’s equiluminance ratio to provide optimal chromatic control. Adding an attention grabbing smooth pursuit target could aid in keeping infant’s engaged during pupillary frequency tagging.

In the future, PFTM may be used by technology companies marketing TrueTone displays on their devices. With the continued refinement of the technology, it may be possible to find a user’s exact equiluminance point with a simple test of less than a minute and then build this luminance ratio into the display algorithms of the device. A screen that is personalised to the colour perception of each user will allow for a more pleasant user experience. This could be especially helpful to colour-blind users, for whom the display can be programmed for optimal contrast based on their unique equiluminance values.

## 4. Methods

### 4.1. Human Participants

The experiments on the three methods of measuring equiluminance were run on 8 participants (5 male and 3 female), including two of the authors. All participants gave informed consent in writing prior to participation and the protocols for the study were approved by the ethical committee of the Institute of Neuroscience, Chinese Academy of Sciences, in accordance with the Declaration of Helsinki. They all reported normal colour vision and normal or corrected-to-normal acuity.

### 4.2. Non-human Participants

Four adult rhesus macaques (male Macaca mulatta, 6 to 8 years old, weighing 4.5 to 10.9 kg) participated in this study. All experimental procedures were approved by the Animal Committee of the Institute of Neuroscience, Chinese Academy of Sciences, and were in accordance with the US National Institutes of Health Guide for the Care and Use of Laboratory Animals.

### 4.3. Apparatus

Stimuli were presented on a linearised Display++ (Cambridge Research Systems, Cambridge, UK) at a frame rate of 120 Hz, a resolution of 1920 x 1080 pixels, and a distance of 67 cm (human, yielding a size of 60.8° × 34.2°) or 72 cm (non-human, yielding a size of 56.9° × 31.8°). We measured 40 samples across the full range of luminance output and CIE spectral characteristics for the red, green and blue channels of the display using a SpectroCal2 spectrophotometer (Cambridge Research Systems, Cambridge, UK). All experiment code and stimulus generation was written using Opticka^54^ and the Psychophysics toolbox (PTB, V3.0.17)^55^, running under MATLAB 2021a & Ubuntu Linux 20.04. All stimuli used 32-bit floating point calculations internally and 14-bits per channel of colour output via the Color++ mode of the Display++. Eye position and pupil diameter were measured from the right eye using an Eyelink 1000 (SR Research, Ontario, Canada) at a sample rate of 1000 Hz.

The Eyelink records the size of the pupil in arbitrary units. We followed the instructions supplied by SR research to generate a scaling factor for a known physical diameter in millimetres. We performed this for both the human and NHP setups separately. Knowing the distance between the infrared camera and the participant, we used an 8 mm black target as an artificial pupil at the same position as the participant’s eye to record the arbitrary units for this known diameter target. The recorded data from the artificial pupil was then used to calculate a scaling factor,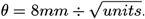. We then used θ to calculate our participant’s pupil diameter using: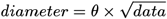.

To ensure precise temporal fidelity of stimulus presentation, we used a photodiode recorded via a LabJack T4 to confirm that temporal intervals were accurate and without significant variance. We also used PTB’s f l i p command to log every frame’s VBL timestamp and calculate any missed frames, to confirm no frames were dropped. The MATLAB code, including friendly GUIs to run experiments for minimum motion, minimum flicker and PFTM methods, as well as analyse the data, is available at <github.com/iandol/equiluminance>.

### 4.4. Minimum Flicker

We used a 15° square of grating presented against a grey background (physically equiluminant with the fixed luminance). Subjects were asked to fixate a small central cross, and maintain fixation within a 2° radius during stimulus presentation, but as this was a matching task we did not enforce fixation. A red/green square-wave grating of 0.25 cycles/° counterphase flickered (red and green exchanged places) at 15 Hz (30 reversals per second). The red and green were the isolated red and green of the display monitor (red and green CIE x / y coordinates were 0.172 / 0.738 and 0.675 / 0.323, respectively) and varied only in intensity. The stimulus profile was a square wave in space and time^33^. The red intensity was fixed to 21.04 cd/m^2^, and the green intensity varied to find the minimum flicker point. The maximum green luminance was 103 cd/m^2^. Before the main trials began, participants were allowed to practice the tasks to become familiar with how their perception of the stimuli changed as they reached their equiluminance point. Participants used key presses to increase or decrease the green luminance of the stimuli. The green luminance value was recorded when the participants achieved minimum flicker. The green value was then randomised, and the same procedure was repeated. Each participant performed five such repeats.

### 4.5. Minimum Motion

We used a 15° square of grating presented against a grey background (physically equiluminant with the fixed luminance). Subjects were asked to initiate fixation on a small central cross for 0.3 seconds and then maintain fixation within a 2° radius during stimulus presentation; failure to maintain fixation aborted that trial. Four repeating frames generated motion to the left for red darker than green and in the opposite direction for red lighter than green (**Figure 5b**). Subjects used the leftward or rightward arrows to choose the direction in a 2-alternate forced choice. When the luminances were matched for the participant, the motion was nulled, and choices were random around 50%. As with the minimum flicker, the minimum motion stimulus had a fixed red luminance of 21.04 cd/m^2^. The yellow was always the mixture of the red and green of the red/green grating at half amplitude ((*R* + *G*) ÷ 2) to keep the mean luminance matched between the red/green and yellow gratings. This was confirmed using our spectrophotometer. The light yellow was 20% more luminous than this (*R* + *G*) ÷ 2 mixture and the dark yellow 20% less luminous. A method-of-constant-stimuli paradigm with 10 repeats was utilised, and psychometric fitting was performed using maximum likelihood estimation with a Quick function (**Equation 1** using the Palamedes toolbox^56^ and MAT-LAB 2021a):

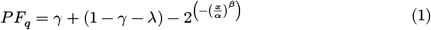

The Quick function estimates the threshold at 50% of the psychometric curve.

### 4.6. Pupil Frequency-Tagging Method

Each trial was initiated by the participant voluntarily foveating (2° radius) a fixation cross^57^ presented against a grey background (physically equiluminant with the fixed value colour). After the initial fixation was maintained for 0.8 to 1.0 seconds, a coloured disc 20° in diameter was presented while the participant continued to maintain fixation for a further 3.5 seconds. The edges of the disc were smoothed using 0.3° hermite interpolation window (OpenGL GLSL `smoothstep` function). Single-colour tasks (where the fixed and variable parameters had the same colour) were used to confirm that the paradigm was effective. In this task there is one step in the middle of the range in which the circle and background are identical, resulting in a solid screen that should produce no pupil oscillations in the tagging frequency band. For measurement of perceptual equiluminance, the luminance of one colour (red) was fixed and the luminance of a second colour (green) chosen from five to twenty linearly spaced luminance steps centered around the physical luminance value of the fixed colour. The duration of each colour presentation in the alternating sequence was 0.283 seconds, yielding a 1.77 Hz tagging frequency (**Figure 1**). The luminance for red was usually fixed at 21.04 cd/m^2^. The grey background was displayed full-screen for 2 seconds between trials to stabilise the pupil diameter before the subsequent trial initiated. Blocks of trials consisted of the same fixed colour which alternated with the variable colour in a fully randomised order with at least 5 repeated presentations of each variable luminance condition.

For analysis, the pupil diameter time series was time-locked to stimulus onset. We first used a 30 ms gaussian smoothing function on the raw pupil data (`smoothdata` in MATLAB). Two different methods were then employed for baseline correction. The first involved calculating the mean pupil diameter between −0.1 ↔ +0.1 seconds for each trial and subtracting it from that trial’s data trace (pre-stimulus baseline correction). The second involved calculating the mean diameter from the same time range used for the subsequent FFT for each trial and subtracting it from that trial’s data trace (mean-baseline correction). Either of these were performed before averaging the trials for each condition. In general, mean-baseline correction resulted in slightly better signal:noise when calculating the FFT, but the differences were minimal (our analysis code computed both to check any differences manually). Next, a discrete Fourier transform was computed from the averaged time-locked signal. To measure the FFT amplitude at the 1st harmonic, the time window must be a multiple of the frequency (e.g. 0.567 × *ρ*, where ρ is the number of repeat oscillation one wishes to measure). We offset the starting time-point manually to where the oscillation was stable (given the onset lag) and the phase of the darkest and lightest variable luminances were diverging so that measured phase was relative to that point. We computed the mean and first harmonic at the tagged frequency and calculated the signal’s phase at the tagged frequency across all variable luminance steps. This was used to compute an interpolated tuning curve fit using a smoothing spline (f i t function in MATLAB), whose minimum is equal to the minimum FFT amplitude at the tagging frequency. We utilised the phase transition to confirm the minimum value. At this minimum, the changing colour is considered the closest perceptual match to the fixed colour.

### Pupil Frequency-Tagging Method applied to non-human primates

to assess PFTM in non-human primates, the paradigm detailed above was adapted to give positive rewards (preferred fruit juice delivered via a peristaltic pump controlled via an Arduino) after successfully initiating and maintaining fixation during the trial. All four non-human primate participants had been previously well trained to fixate on a central fixation spot (a standard procedure for most non-human primate research involving visual stimulation). The paradigm used the same spatial and temporal stimulus structure as that used with the human participants: a central disc and background in which a fixed and variable colour reversed lasting 3.5 seconds in each trial (except for one participant who could only fixate for 3 seconds). The luminance for red was usually fixed at 20.24 cd/m^2^ as we used a different Display++ monitor with a slightly different overall luminance. Given that non-human participants worked for reward, we could test more steps of the variable luminance and more repeated trials for each luminance could be collected. The analysis of the pupil signals was identical to the human data.

### 4.7. Statistical Analysis

Statistical analysis was performed with JASP^58^. We checked for bivariate normality of the correlation using the Shapiro-Wilk test, equality of variance using Levene’s test, and the post-hoc analysis of the repeated-measures ANOVA was corrected using Holm’s method. Group comparisons were visualised using the Raincloud plot library in R^59^.

